# Diversity of exopolysaccharide cluster I in the *Bradyrhizobium* genus

**DOI:** 10.1101/2025.07.22.666213

**Authors:** Sachiko Masuda, Ken Shirasu, Yasuyuki Kawaharada

## Abstract

*Bradyrhizobium*, the largest rhizobial genus, is characterized by variety exopolysaccharides (EPSs) components, depending on the species. However, several genes involved in EPS synthesis remain unknown. In this study, we investigated whether 186 *Bradyrhizobium* strains possess EPS cluster I, which is involved in the synthesis of a pentasaccharide EPS by *B. diazoefficiense* USDA110. Homologous genes involved in EPS synthesis were found to be absent in the *B. elkanii* and Photosynthetic *Bradyrhizobium* supergroups, other than the *B. japonicus* supergroup. These findings suggest that genes related to EPS synthesis are yet to be identified in the *B. elkanii* and Photosynthetic *Bradyrhizobium* supergroups.

## Main text

Exopolysaccharides (EPSs), produced by soil bacteria, protect the bacteria from various environmental stresses (Srivastava and Sharma, 2023). In the context of plant-microbe interactions, bacterial EPSs protect against reactive oxygen species, boost plant defense responses, and promote adhesion to the host plant root (Morcillo and Manzanera, 2021). Moreover, low molecular weight EPSs (LMW EPSs) produced by nodulating bacteria, known as rhizobia, serve as signaling molecules during nodule formation in legumes (Brett J. Pellock *et al*., 2000). These LMW EPSs are recognized by the LysM-type receptor exopolysaccharide receptor 3 (EPR3), which regulates infection progression to the host cells (Kawaharada *et al*., 2015, 2017).

*Bradyrhizobium* spp., which form symbiotic nodulation with major legume crops, produce structurally distinct EPSs compared to other rhizobial genera, such as *Ensifer, Rhizobium*, and *Mesorhizobium* (Acosta-jurado *et al*., 2021). The EPS secreted by *B. diazoefficiens* USDA110 is a pentasaccharide composed of D-mannose, 4-O-D-methylgalactose, D-glucose and D-galacturonic acid with an acetyl group substitution (Louch and Miller, 2001; Mort and Bauer, 1982). Similarly, *B. diazoefficiens* USDA123 secretes an EPS of comparable composition, whereas within soybean nodules, the differentiated bacteroids secrete an EPS mainly composed of rhamnose (known as nodule polysaccharides: NPS) (An *et al*., 1995; Huber *et al*., 1984). In contrast, *B. elkanii* USDA31, USDA39, and USDA76 produce tetrasacccharide EPS and NPS composed of 4-O-D-methylgalactose and D-rhamnose (An *et al*., 1995; Huber *et al*., 1984; Streeter *et al*., 1992).

In *B. diazoefficiens* USDA110, the genes responsible for EPS synthesis are organized into two EPS clusters on the bacterial chromosome: EPS cluster I (containing *exoU, exoM, metA, exoP, exoT*, and *exoB*), and EPS cluster II (containing *lspL, ugdH, blr2358*, and putative *exoP* and *exoZ*) (Becker *et al*., 1998; Kaneko *et al*., 2002; Parniske, 1993; Quelas *et al*., 2010; Xu *et al*., 2021). Notably, the strain containing the mutant *exoB*, which encodes UDP-galactose 4-epimerase, failed to produce EPS, resulting in dry colonies compared to the wet colonies observed in the wild-type on the solid medium (Quelas *et al*., 2010). Three independent research groups investigated the nodulation phenotype of *exoB* mutants in soybeans. While Quelas et al. reported that the USDA110 *exoB* mutant is a nodule-defective phenotype, Parniske et al. and Becker et al. observed that the *B. diazoefficiens* USDA110spc4 *exoB* mutants formed delayed and fewer nodules. Furthermore, when these mutants were treated with the EPSs produced by *B. diazoefficiens* USDA110spc4, nodule number restored (Becker *et al*., 1998; Parniske, 1993; Quelas *et al*., 2010). *Ensifer, Rhizobium*, and *Mesorhizobium* produce octasaccharide EPSs. The genes involved in biosynthesis, polymerization, and excretion have been identified (Acosta-jurado *et al*., 2021). In contrast, among the *Bradyrhizobium* genus, although the different EPS components in *B. diazoefficiens* and *B. elkanii* have been characterized, the genes related to their synthesis have only been identified in *B. diazoefficiens* USDA110. In this study, we investigated the distribution of genes essential for EPS synthesis within EPS cluster I of *B. diazoefficiens* USDA110, a member of the *Bradyrhizobium* genus, using a *Bradyrhizobium* pangenome.

We used of the complete genomic sequences from 187 *Bradyrhizobium* species (**Supplemental Table S1**). These sequences were obtained from the National Center for Biotechnology Information (NCBI) database with completeness ≥ 90, contamination ≤ 5 and checkm quality ≥ 50 based on the GTDB-tk database (bac120_metadata.tsv, released on 2025-04-08). Some genomes retrieved from GTDB-Tk were suppressed in the NCBI database due to contamination, while duplicate genome submissions for the same strain were found in other cases. These contaminating and duplicate sequences were removed. The complete genomes of 187 *Bradyrhizobium* strains and *Azospirillum* sp. B510 (Elbeltagy *et al*., 2001) as an outgroup were used to construct a maximum-likelihood phylogenetic tree inferred using GTDB-Tk v2.4.1 with the de_novo_wf workflow (Chaumeil *et al*., 2022), IQ-TREE v2.4.0 (Minh *et al*., 2020), and visualized with FigTree v1.4.4 (http://tree.bio.ed.ac.uk/software/figtree/). The homologous genes related to the surrounding EPS cluster I were searched using DIAMOND (v 2.1.10) (Buchfink *et al*., 2021) with BLASTp using the *B. diazoefficiens* USDA110 genes as query sequences. The threshold was set at ≥70% identity or E-value ≤1e-05. When multiple homologous genes were dispersed throughout the genome, only those located on or near the EPS cluster I, if present, were included in the gene map. In the absence of EPS cluster I, only the homologous gene with the lowest E-value among the dispersed candidates was included in the gene map. Gene maps were created using the ‘gggenes’ package in R (https://www.R-project.org/).

Phylogenetic analysis based on whole genome sequences classified 187 *Bradyrhizobium* strains into three supergroups: (I) *B. japonicum*, (II) *B. elkanii*, and (III) Photosynthetic *Bradyrhizobium* (PB), consistent with previous reports (**Figure 1**) (Avontuur *et al*., 2019; Camuel *et al*., 2023). To investigate the distribution of genes within EPS cluster I across the *Bradyrhizobium* genus, we searched for homologs of 17 genes surrounding this cluster in 186 *Bradyrhizobium* genomes. Analysis using BLASTp revealed that homologous genes outside EPS cluster I, such as *blr7567, blr7568, blr7580*, and *blr7582*, were conserved across most *Bradyrhizobium*, regardless of supergroup classification. In contrast, multiple homologous genes inside the cluster-particularly *exoU* and *exoM*, were absent in most strains from the *B. elkanii* and PB supergroups (**Figure 1, Supplementary Table 2**). When the *B. elkanii* supergroup was further divided into six subgroups, subgroups 2, 3, and 5, as well as the PB supergroup, lacked most of the homologous genes in the EPS cluster I (**Figure 1**). To further understand these homologous genes, we conducted a comparative analysis using representative *Bradyrhizobium* from each supergroup (**Figure 2**). *B. japonicum* USDA6 and *B. ottawaense* SG09 in the *B. japonicum* supergroup were found to retain the *exoU, exoM, metA, exoT*, and *exoB* homologous genes in their cluster. While *B. elkanii* USDA76, *B. septentrionale* 1S1, and *Bradyrhizobium* sp. ORS278 in the *B. elkanii* and PB supergroups, respectively, possessed various gene compartments. In *B. elkanii* USDA76 in the *B. elkanii* supergroup-subgroup 2, the homologous genes located outside the EPS cluster I (e.g., *blr7566, blr7567, blr7568, bll7589, blr7581*, and *blr7582*) were conserved, whereas the genes within the cluster were replaced with unknown genes (**Figure 2**). In contrast, in *B. septentrionale* 1S1 in the *B. elkanii* supergroup-subgroup 3, several homologous genes were identified (**Figure 1**), but these genes did not from any clusters (**Figure 2**). In *Bradyrhizobium* sp. ORS278 in the PB supergroup, the genes were scattered across different genomic regions (**Figure 2**).

**Figure 1.**
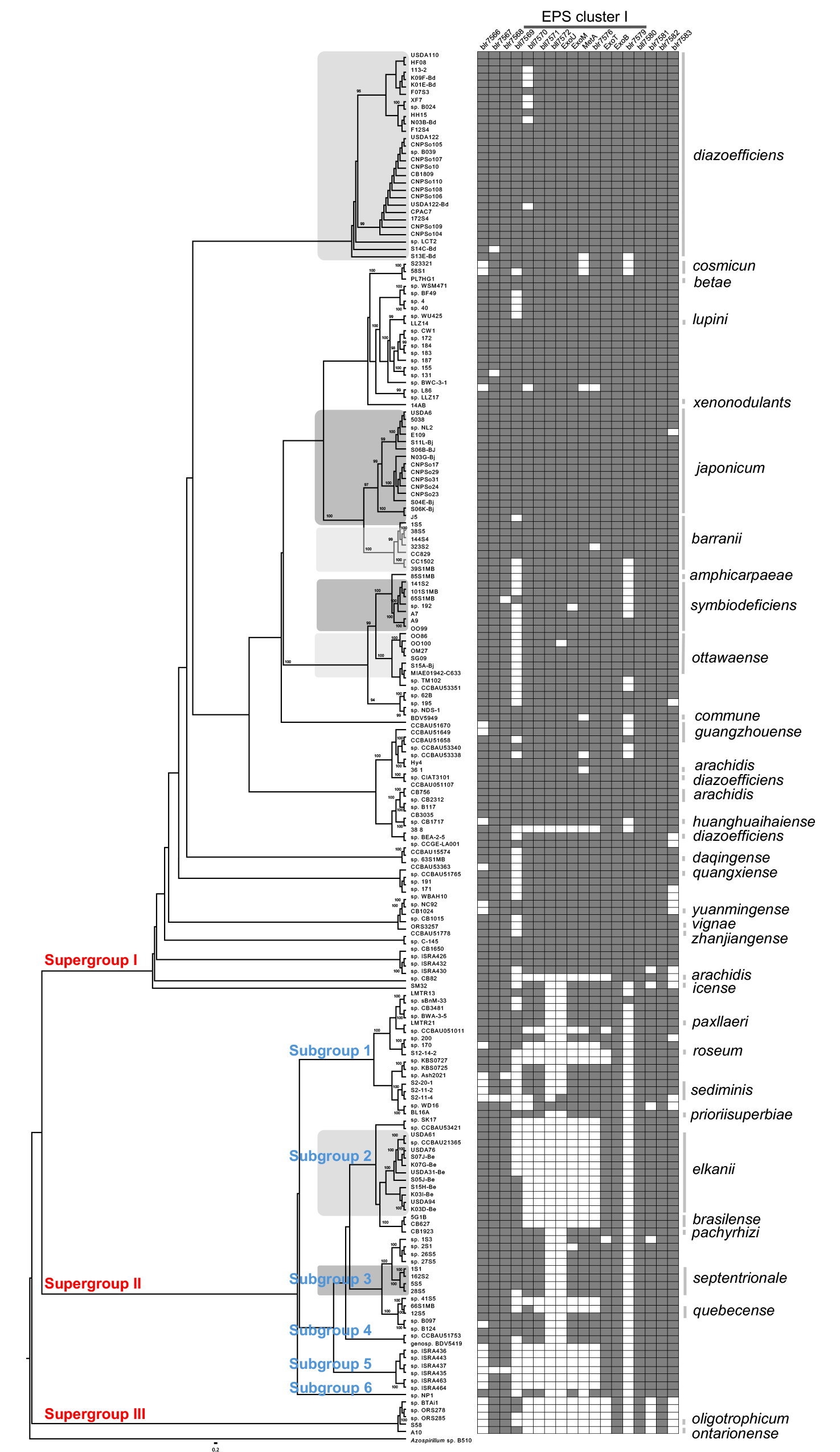
Phylogenetic tree of the *Bradyrhizobium* genus based on the genes related to EPS synthesis. A maximum-likelihood phylogenetic tree based on whole genomes from 187 *Bradyrhizobium* strains and *Azorhizobium* sp. B510 as an outgroup. Bootstrap values were 1000 replicates. The substitution rate was estimated at 0.2 per nucleotide position. The boxes on the right indicate homologous genes surrounding EPS cluster I of *B. diazoefficiens* USDA110 in each *Bradyrhizobium* strain. Gray boxes represent genes with more than 70% amino acid sequence identity or an E-value of less than 1e-05. Each gene ID is shown in **Supplemental Table 2**.

**Figure 2.**
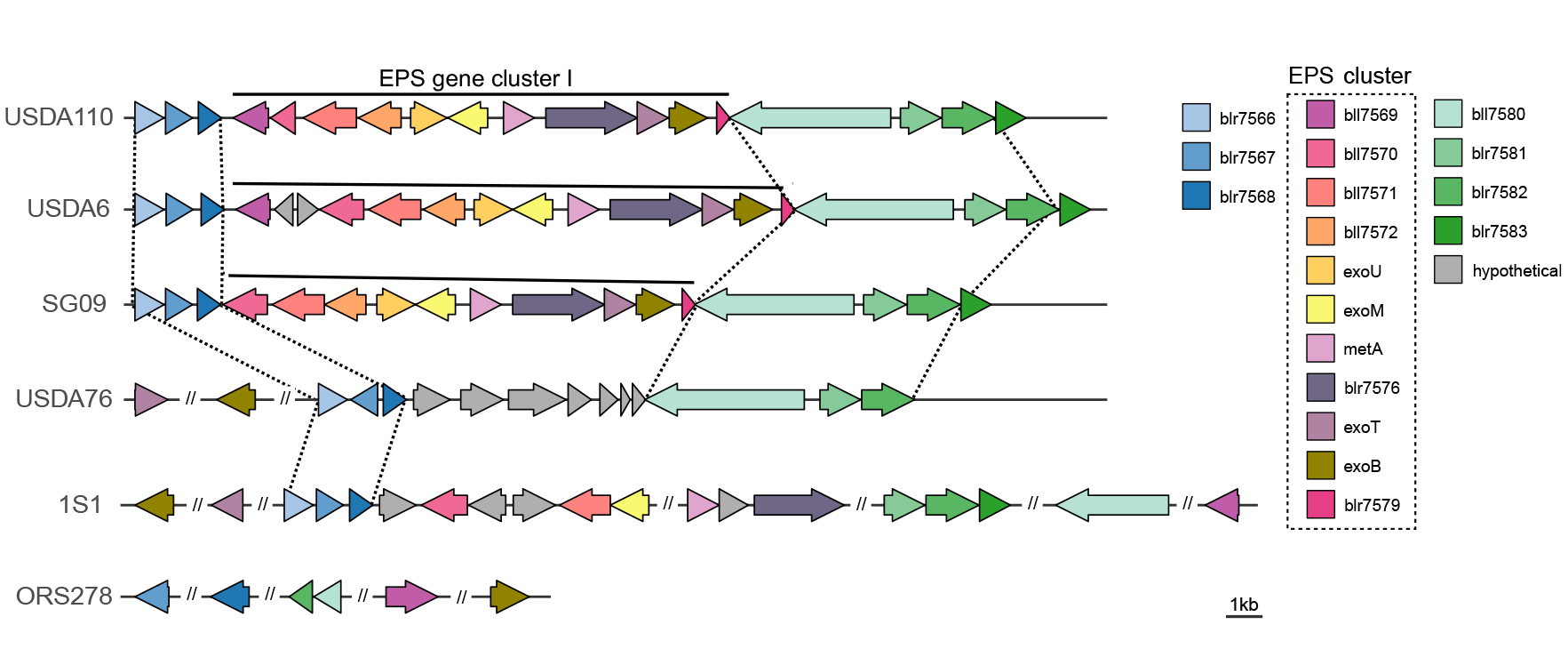
Comparison of EPS cluster I alignment in representative *Bradyrhizobium* strains. Distribution of the homologous genes in EPS cluster I in. *B. japonicum* USDA6 and *B. ottawaense* SG09 (*B. japonicum* supergroup), *B. elkanii* USDA76 (*B. elkanii* supergroup-subgroup 2), *B. septentrionale* 1S1 (*B. elkanii* supergroup-subgroup 3), and *Bradyrhizobium* sp. ORS278 (PB supergroup) using the sequences surrounding EPS cluster I of *B. diazoefficiens* USDA110 as queries.

Based on these results, we concluded that the EPS in the *B. japonicum* supergroup, including *B. diazoefficiens* USDA110, is a pentasaccharide (Louch and Miller, 2001; Mort and Bauer, 1982), and were identified the genes in EPS cluster I (Becker *et al*., 1998; Parniske, 1993; Quelas *et al*., 2010). While *B. elkanii*, unlike *B. diazoefficiens* USDA110, produced a tetrasaccharide EPS (An *et al*., 1995; Huber *et al*., 1984; Streeter *et al*., 1992), the genes responsible for their synthesis were missing among the homologous genes of *B. diazoefficiens* USDA110. These findings suggested that the genes responsible for EPS synthesis in the *B. elkanii* and PB supergroups have yet to be identified.

## Acknowledgements

This work was supported by New Energy and Industrial Technology Development Organization (JPNP18016) to K.S and Y.K. RIKEN-TRIP program to K.S.

## Author contribution

S.M and Y.K designed the study. S.M and K.S performed bioinformatics analysis, and S.M and Y.K wrote all manuscripts.

## Supplementary information

**Supplemental Table S1. All data for the genome ID in this study**.

**Supplemental Table S2. Homologous genes surrounding the EPS cluster I in each *Bradyrhizobium* strain**.

